# Emergent Patterns from Probabilistic Generalisations of Lateral Activation and Inhibition

**DOI:** 10.1101/033951

**Authors:** Lisa Willis, Alexandre Kabla

**Affiliations:** The Sainsbury Laboratory, University of Cambridge, Cambridge CB2 1LR, UK; Department of Engineering, University of Cambridge, Cambridge CB2 1PZ, UK

**Keywords:** lateral inhibition, pattern formation, nonlinear stochastic model

## Abstract

The combination of laterally activating and inhibiting feedbacks is well known to spontaneously generate spatial organisation. It was introduced by Gierer and Meinhardt as an extension of Turing’s great insight, that two reacting and diffusing chemicals can spontaneously drive spatial morphogenesis *per se*. In this study, we develop an accessible nonlinear and discrete probabilistic model to study simple generalisations of lateral activation and inhibition. By doing so, we identify novel modes of morphogenesis beyond the familiar Turing-type modes; notably, beyond stripes, hexagonal nets, pores, and labyrinths, we identify labyrinthine highways, Kagome lattices, gyrating labyrinths, and multi-colour travelling waves and spirals. The results are discussed within the context of Turing’s original motivating interest: the mechanisms which underpin the morphogenesis of living organisms.

## 1 Introduction

As a complex multicellular organism grows and develops, each one of its cells follows the same set of genomically encoded instructions, yet different cells beget drastically different fates so bringing about the organism’s complex structure. Turing was among the first to present a powerful idea pertaining to this phenomenon, when he realised that a spatially homogeneous soup of just two chemically reacting species can spontaneously morph into a structured pattern owing to nothing more than the diffusion of these species, so long as the reaction kinetics are of the appropriate sign and the diffusivities are sufficiently different [1]. Later Gierer and Meinhardt extended this notion to show how processes other than reaction-diffusion can potentially drive pattern formation: they demonstrated that any process which feeds back on itself over two lateral ranges—one short range that quickens or activates the process, the other long range that competitively slows or inhibits it—can spontaneously generate structural organisation [2, 3]. This combination of feedbacks has come to be known as short range activation and long range inhibition and is widely accepted as the key criteria for Turing-type patterning [3, 4, 5].

Groups of cells can effect lateral activation/inhibition by, for example, secreting ligands that diffuse on average a few cell lengths before degradation or binding to receptors; this binding triggers an intra-cellular signalling cascade which in turn increases/decreases the ligand’s expression. Given the vast number of intra- and inter-cellular signals operating within and between cells, it seems probable that other types of lateral feedback beside lateral activation and inhibition also operate during development; furthermore, the structural diversity among living organisms is immense whereas lateral activation and inhibition alone generate a limited range of patterns. What other types of lateral feedback may be operating to bring about the development of a multicellular organism? What patterns can be generated by recombinations of these feedbacks and what feedbacks are necessary and sufficient to generate a particular pattern?

Here, we study a model of interacting lattice sites that laterally activate and inhibit one another [6]. The lattice sites may be considered as a layer of immobile discrete cells, each executing a dynamic differentiation program to transit between boolean states according to a common set of deterministic instructions but subject to some level of noise. The model has an important distinguishing feature: its discrete and probabilistic formulation makes it straightforward to introduce and to simulate new pattern-generating modules—new classes of lateral feedback operating alongside lateral activation and inhibition—in a systematic way: the discreteness allows us to write down simple equations governing the bifurcation diagrams, while the noise provides an in-built test of robustness against fluctuations and renders the long-term patterning dynamics independent of the model’s initial state. For lateral activation and inhibition only, the model’s final states of morphogenesis are either striped, hexagonally netted, labyrinthine, spotted, or uniform block colour, patterns that are ubiquitous in nature and are well known to be generated by Turing-type models [6]; whereas the new feedback modules that we introduce generate a number of surprising static and dynamic patterning modes that are, to the best of our knowledge, new. The ubiquity in nature of the patterning modes generated by the original model—stripes, hexagons, labyrinths, spots—(see e.g. [7, 8]; reaction-diffusion [1, 9, 10, 11]; directional solidification [12]; granular/fluid flows [13], hydrodynamic instabilities, animal furs, seashells, and more [7, 14, 15]) suggest that concrete applications for these novel patterning modes may soon be discovered.

## 2 Model and Methods

### 2.1 Model

The core model of discrete and probabilistic lateral activation and inhibition introduced in [6] is now described. In each section of the results, this core model is generalised in a different way.

The model runs on a 2D square grid of lattice sites that are either black (*B*) or white (*W*). From an initial configuration, lattice sites flip their colour, from black to white or white to black, one at a time. The rate at which a particular site flips its colour is determined by the following three rules:

i. a site can flip to a colour only if a neighbouring site has that colour (this is *activation at the interface*);
ii. the likelihood of flipping to a colour decreases with the density of sites of that colour within a particular long range *r_L_* (this is *long-range inhibition*), where *r_L_* ≫ 1 is measured in units of a lattice site diameter;
iii. the likelihood of flipping to a colour increases with the density of sites of that colour within a particular short range *r_S_* (this is *short-range activation*).

A noise level *T_L_* is associated with the long-range inhibition: as *T_L_* increases, the long-range inhibition is wiped out. Similarly, a noise level *T_S_* is associated with the short-range activation. A parameter, *β*, determines the strength of the propensity for cells to flip to a particular colour, either black or white, independently of the lateral activation and inhibition. When *β* = 0, this propensity is zero, then colour configurations converge to attractors that are 50% white and 50% black on average over multiple instances of the simulation; in this case we say there is *colour symmetry* and the model’s specification is unchanged when white is interchanged with black (see the reaction kinetics (1) below). Perturbing *β* away from zero, then rather colour configurations accumulate an excess of one particular colour—white is more abundant for *β* > 0, while black is more abundant for *β* < 0—and so *β* is called a *symmetry breaking parameter*, see Figure 1a.

**Figure 1:**
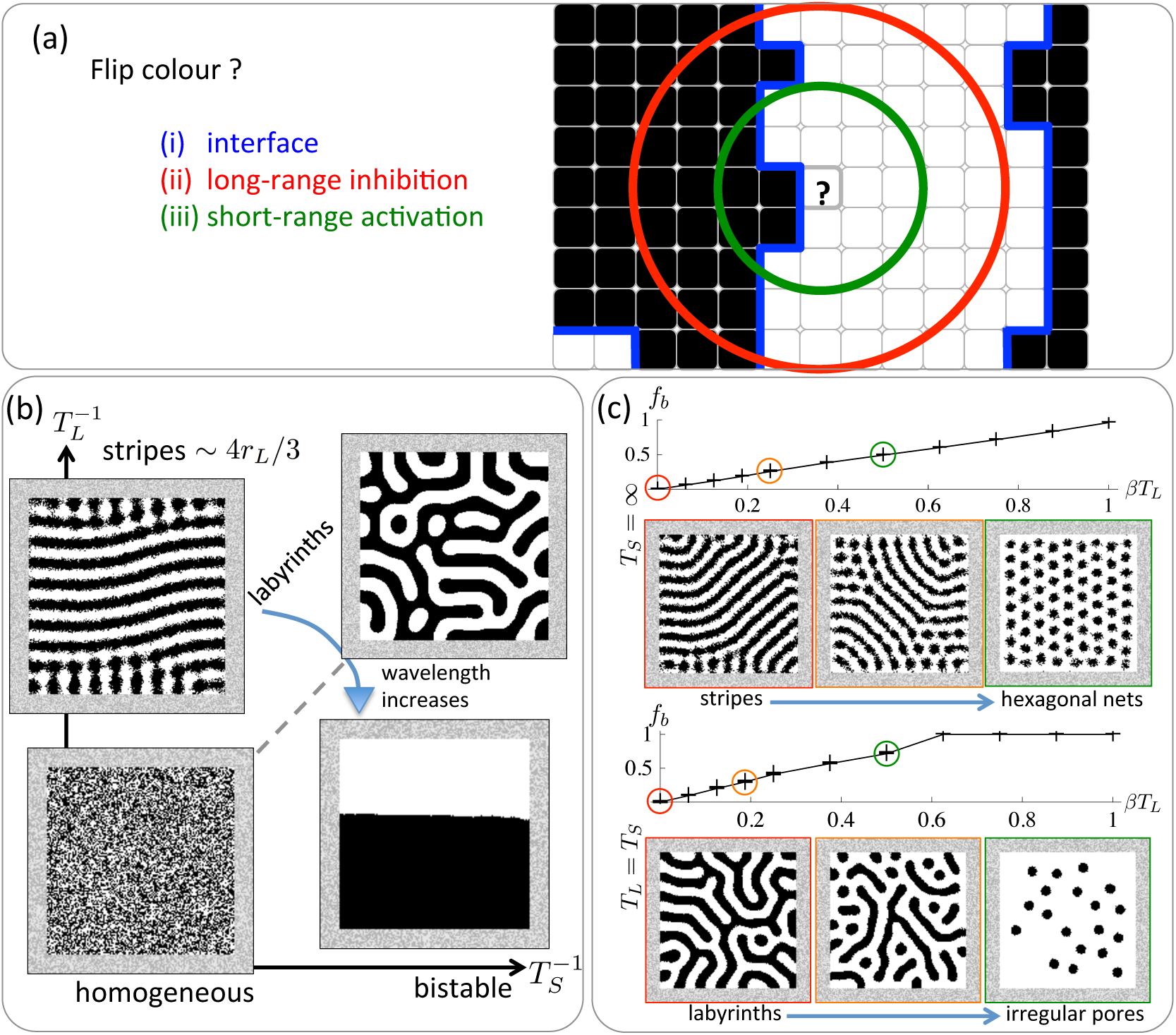
A probabilistic and discrete model of lateral activation and inhibition. (a) The model has three key rules: (i) activation at the interface; (ii) long-range inhibition; and (iii) short-range activation, (b) Sketch bifurcation diagram for colour symmetry (*β* = 0): for lateral inhibition only (*T_L_* ≪ 1, *T_S_* = ∞), dynamics converge to stationary stripes; for short-range activation and long-range inhibition together (*T_L_* ≈*T_S_* ≪ 1), stripes bifurcate to stationary labyrinths; weakening the long-range inhibition (*T_L_* ≫ *T_S_*), eventually labyrinths bifurcate to bistable attractors, (c) For colour symmetry breaking (*β* ≠ 0), stripes transit to hexagonal nets while labyrinths transit to irregular arrangements of pores; these transitions are represented by plots of *f_b_*, the summary statistic for 2-colour symmetry breaking transitions (upper plot is for lateral inhibition only; lower plot is for short-range activation and long-range inhibition together). The correspondence between patterns and *f_b_* is colour coded by red/orange/green. Simulation parameters and numbers of instances are listed in Table 1.

Precisely, from a given colour configuration, the probability that the next colour flip is at site x to colour *C* = *W* or *B* is non-zero if and only if any of the 8 neighbours of x has colour *C* (the set of neighbouring colours is denoted ℕ_x_); this probability has one of two possible forms depending on the colour *C_x_* of site x

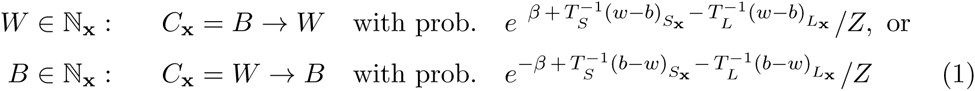

where 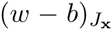 is the fraction of white minus black sites within range *r_J_* around x, and *Z* is the colour configuration-dependent normalising factor, or equivalently the sum of numerators in (1) over every non-zero probability colour flip. This completely specifies the model’s reaction kinetics, except for initial and boundary conditions which are described in the next section. For the corresponding continuous time definition and a partial derivation of the mean-field equations see Supplementary Material.

When the model is run, invariably it converges to a macroscopically stationary attractor that is either patterned, or homogeneously noisy, or uniform block colour (see Figure 1b and 1c and Movies S1 and S2 and [6]). Patterned attractors are time-invariant or stationary if and only if sites along their interfaces flip from black to white and back again from white to black at equal rates, so from (1) we have the following necessary and sufficient mean-field approximation for stationary attractors,

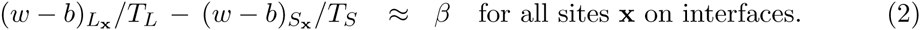

For lateral inhibition only and no symmetry break (high noise on the short-range *T_S_* = ∞, low noise of the long-range *T_L_* ≪ 1, and *β* = 0), from (2) stationary attractors have 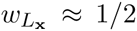 along interfaces (because 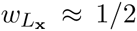), whereas for both lateral activation and inhibition with no symmetry break (*T_S_* = *T_L_* ≪ 1 and *β* = 0) then we have 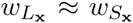 along interfaces. Straight interfaces satisfy the condition 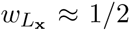, whereas 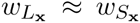 can be satisfied by curved interfaces. Moreover, the mean-field approximation predicts a linear increase of 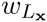 with *β* for lateral inhibition only (*T_S_* = ∞ in (2)).

### 2.2 Methods

Each model was simulated on a 2D square grid of (*l* + 2*r_L_*) × (*l* + 2*r_L_*) sites. Initially, all sites of the simulation are equally and independently likely to be one colour among a list of permitted colours (black or white in the model above; the list is extended in the generalisations below). Thereafter, sites within the central square domain of *l* × *l* lattice sites flip their colour according to the reaction kinetics specified in each section. Outside of this *l* × *l* domain, colours remain fixed for all time—this constitutes the boundary condition.

Simulations of the core model show that the local solutions predicted by the mean-field approximation (2) are invariably realised: for colour symmetry and lateral inhibition only (*T_S_* = ∞, *T_L_* ≪ 1, *β* = 0), attractors are approximately straight stripes, whereas curved labyrinths are generated by lateral activation and inhibition together (*T_S_* = *T_L_* ≪ 1, *β* = 0), see Figure 1b, Movies S1 and S2, and [6]. Now breaking the colour symmetry by increasing *β* > 0, then stripes bifurcate to white hexagonal nets while the predicted linear increase of 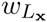 with *β* holds true for the overall fraction of white sites in the *l* × *l* domain, as shown in the plots in Figure 1c which are fully described below. The initial and boundary conditions appear to have no effect on the local structure of the final pattern so long as simulations are run until colour configurations have converged to an attractor [6]. Near to the boundary, stripes and labyrinths tend to align perpendicularly to the boundary and in this way the domain’s geometry influences patterns’ orientations, see Supplementary Material. The wavelength of stripes, and of labyrinths when *r_S_* ≪ *r_L_*, is 4*r_L_*/3 see Supplementary Material and [6]; increasing the domain size *l* × *l* appears to have no effect on this wavelength. In all simulations, the domain size was set such that the wave number *l*/4*r_L_*/3 > 8.

We have seen, then, that in this model the combination of the three rules (i) activation at the interface, (ii) short-range activation, and (iii) long-range inhibition generates a range of Turing-type patterns—stripes, labyrinths, hexagonal nets, and spots—and, moreover, that in several cases the correspondence between stationary patterning modes and parameter values, i.e. the bifurcation diagrams, can be roughly anticipated simply by inspecting the mean-field approximation (2). Therefore, we searched for interesting new attractors by retaining the three rules (i)–(iii) while systematically generalising the model in different simple ways. Importantly, all generalisations in this study introduce new linearly independent terms into the mean-field approximation for stationary interfaces: we hypothesized that only such generalisations can produce attractors with interesting new patterns, see Supplementary Material. We consider each linearly independent term in the mean-field approximation to be a *lateral feedback module*. Each lateral feedback module has a corresponding free parameter.

In order to quantitatively represent transitions between patterning modes, a representative summary statistic was computed from each instance of the simulation and its variation with the corresponding parameter was plotted. White/black colour symmetry breaking transitions, where *β* is varied from zero, are quantitatively represented by the variation in *f_b_* which represents the fractional excess in white over black sites in the *l* × *l* domain, as shown in the plots in Figure 1c. In order to generate these plots, the simulation was run 5 times for each value of *β* with all other parameters held constant. The cross-hairs are *f_b_* computed for each instance of the simulation; the line-graph connects the averages of *f_b_* for different values of *β*. A similar format is followed for every plot in the article. Table 1 lists the corresponding parameter values and the number of repeated instances of the simulation. Other summary statistics are introduced in Results; the corresponding and complete descriptions can be found in Supplementary Material. These summary statistics do not depend on the domain size or the end time of the simulation.

## 3 Results

In all sections, for a clear portrayal of the model’s dynamics *it is essential to view the corresponding movies*. Parameters for each movie are listed in Supplementary Material.

### 3.1 Labyrinthine highways, train tracks, and Kagome lattices from a competing nonlinear inhibition on the short range

A natural extension of the model, which introduces two new lateral feedback modules while retaining rules (i)–(iii), includes in the exponent symmetry breaking nonlinear terms 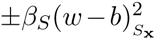 associated with the short range, and symmetry preserving nonlinear inhibitory terms 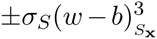 which compete with the short range activation. All other notation is left unchanged. The reaction kinetics are:

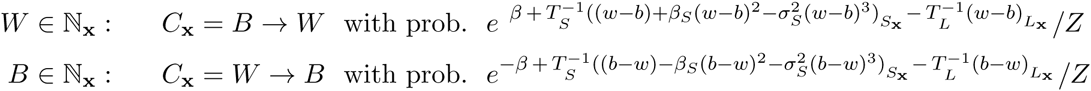

where *Z* is the colour configuration-dependent normalising factor. We focus on perturbations to labyrinth attractors: in all simulations *T_S_* = *T_L_* = 1/16 ≪ 1. Movies S3–S5 and Figure 2 portray the dynamics.

**Figure 2:**
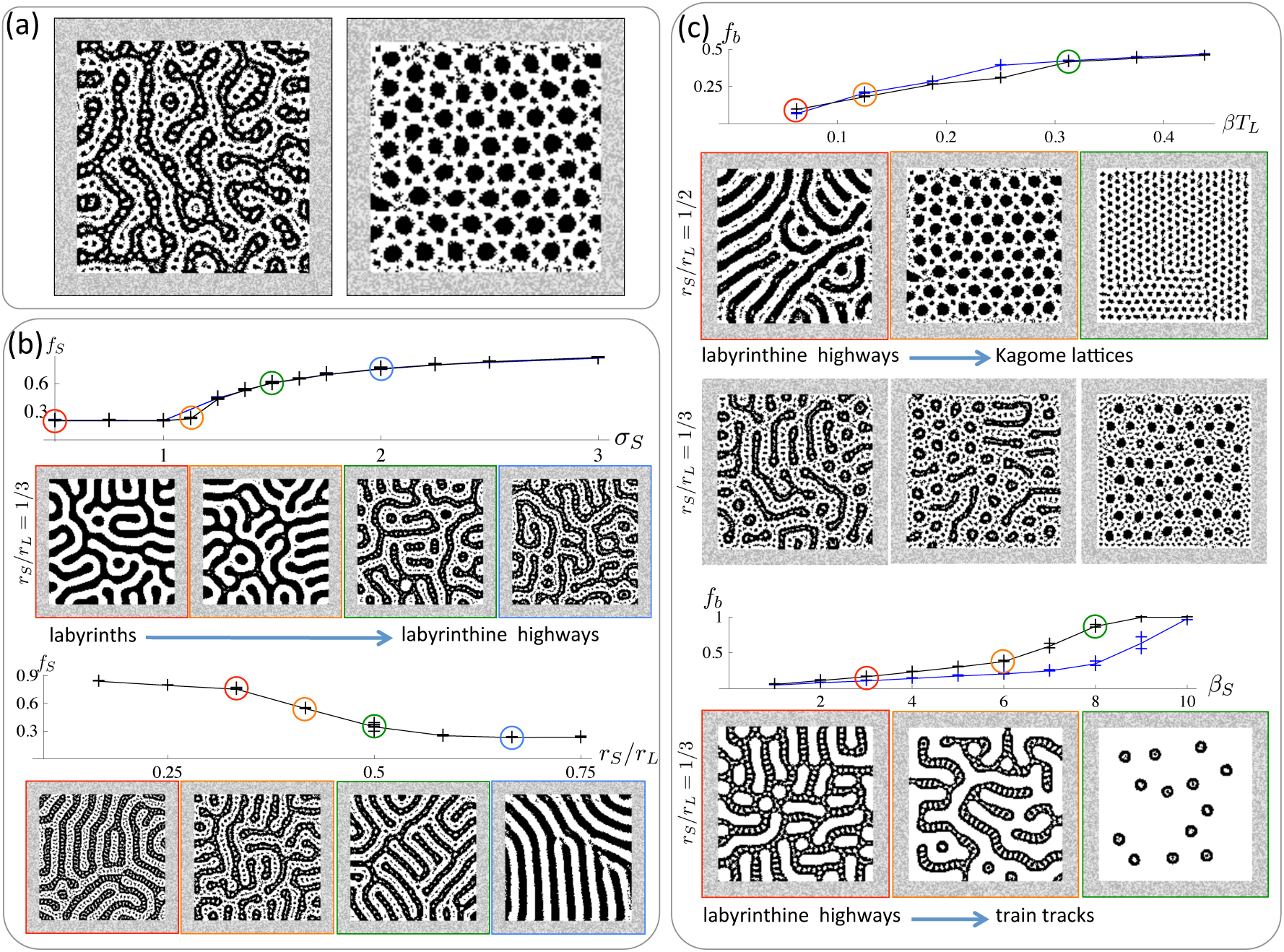
Labyrinthine highways and Kagome lattices for competing nonlinear inhibition on the short range. In all simulations, *T_S_* = *T_L_* = 1/16; other parameters are in Table 1. (a) A nonlinear inhibitory lateral feedback that competes with the activatory feedback on the short range generates labyrinthine highways (left) and Kagome lattices (right), (b) Colour symmetry. Increasing |*σ_S_*| > 1 causes labyrinths to bifurcate to labyrinthine highways as represented by the sudden increase in *f_S_* (the black curve superposed on the blue curve is for *1*, *r_S_* and *r_S_* rescaled by a factor of 1.5 while other parameters are held constant). The bifurcation occurs only when *r_S_*/*r_L_* <1/2 (lower plot), (c) Colour symmetry break. In all plots *σ_S_* = 2. Increasing *β* causes labyrinthine highways to bifurcate smoothly to Kagome lattices for *r_S_*/*r_L_* = 1/2 (upper blue plot) or interwoven short and long-scale hexagonal nets for *r_S_*/*r_L_* = 1/3 (upper black plot). Increasing *β_S_* causes labyrinthine highways to bifurcate to train tracks (lower plots; blue (black) curve is for *r_S_*/*r_L_* = 1/2 (1/3)).

For colour symmetry (*β*,*β_S_* = 0), as depicted in Figure 2b and Movie S3, increasing |*σ_S_*| > 1 causes labyrinths to suddenly bifurcate to near-stationary labyrinthine highways so long as *r_S_*/*r_L_* < 1/2. The summary statistic representing this bifurcation in the plots of Figure 2b, denoted by *f_S_*, measures the prevalence of the short-range spots within the labyrinth tracks, see Supplementary Material for details. Values of *f_S_* appear to be unaltered when *l*, *r_L_*, and *r_S_* are simultaneously rescaled while other parameters are held constant, indicating that discretisation effects of the lattice are small (compare the blue and black curves which are superposed in the upper plot of Figure 2b).

The bifurcation point |*σ_S_*| = 1 can be qualitatively explained as follows. So long as |*σ_S_*| < 1, the net short-range feedback is always activatory for all sites since 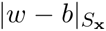 is bounded above by 1—there is no possibility that the short-range feedback switches to inhibitory. When |*σ_S_*| > 1, the net short-range feedback switches sign to become inhibitory for any configuration such that 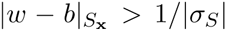, then qualitatively new dynamics are possible.

When colour symmetry is broken by perturbing *β* ≠ 0, for *r_S_*/*r_L_* = 1/2 labyrinthine highways transit to Kagome lattices—a pattern of intermeshed regular hexagons and triangles where the diameter of the hexagon is twice the side of the triangle—that are best known to feature in the atomic arrangement of the minerals Herbertsmithite and jarosite. Whereas for *r_S_*/*r_L_* = 1/3, labyrinthine highways transit to interwoven short and long-scale hexagonal nets (for *r_S_*/*r_L_* = 1/3) as depicted in Figure 2c and Movies S4 and S5. When the short-range symmetry breaking term is perturbed *β_S_* ≠ 0, labyrinthine highways transit to train tracks. As both *β* and *β_S_* are varied, the transitions between patterns are apparently gradual and smooth, as demonstrated by the plots of Figure 2c which show gradual and smooth increases of *f_b_* (the blue curve is for *r_S_*/*r_L_* = 1/2; the black curve is for *r_S_*/*r_L_* = 1/3).

### 3.2 Gyrating labyrinths from a competing nonlinear activation on the long range

A second natural extension of the model, analogous to the extension of Section 3.1, includes in the exponent symmetry breaking nonlinear terms 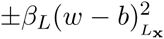 associated with the long range, and symmetry preserving nonlinear activatory terms 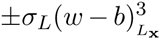 which compete with the long range inhibition. All other notation is left unchanged. The reaction kinetics are:

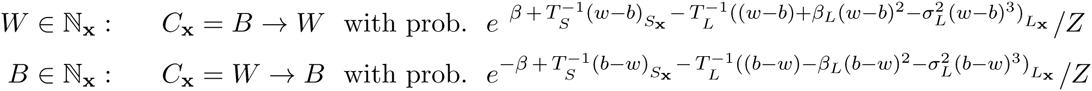

where *Z* is the colour configuration-dependent normalising factor. Again, we focus on perturbations to labyrinths: in all simulations *T_S_* = *T_L_* = 1/16 ≪ 1. Movie S6 and Figure 3 portray the dynamics.

**Figure 3:**
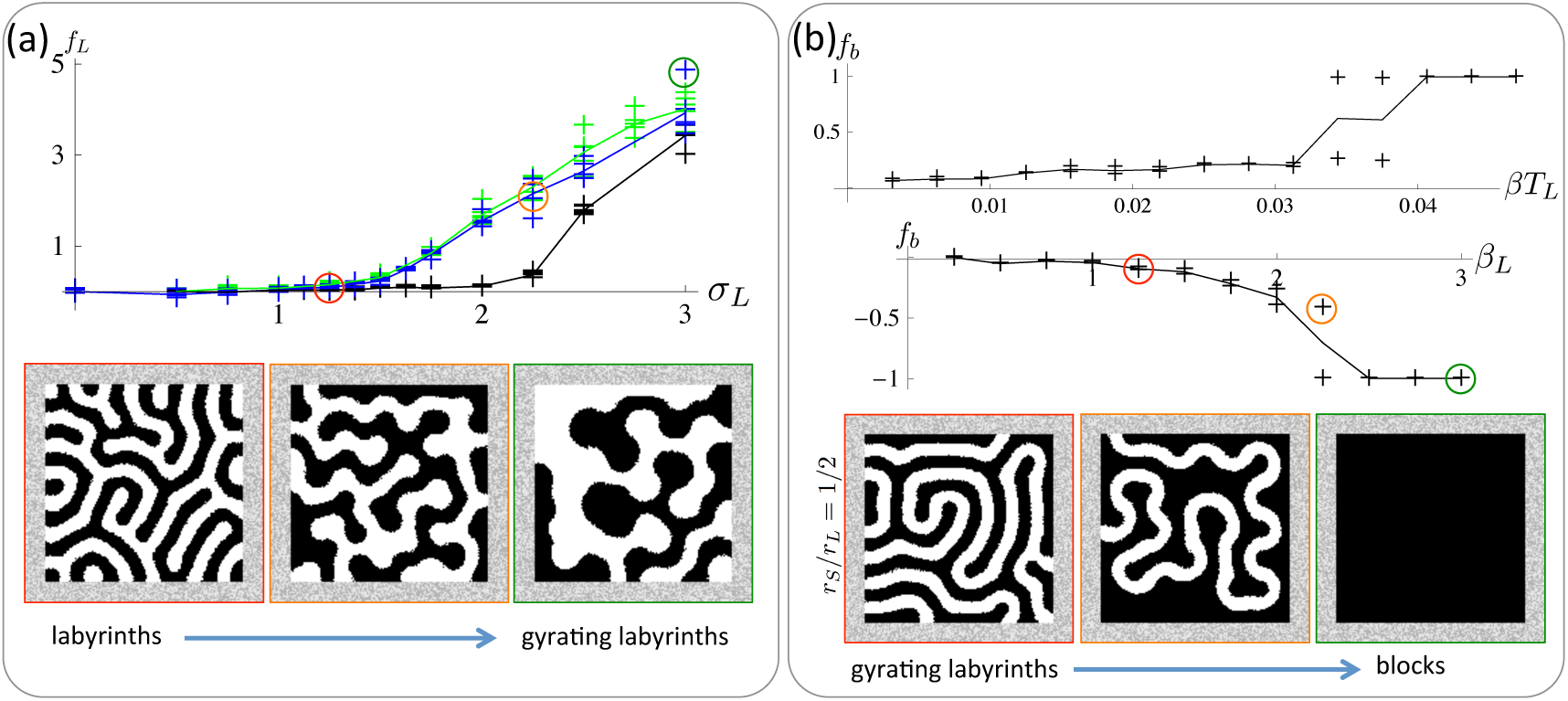
Dynamic gyrating labyrinths for competing nonlinear activation on the long range. In all simulations, *T_S_* = *T_L_* = 1/16; other parameters are in Table 1. (a) Colour symmetry. Labyrinths bifurcate to continually gyrating labyrinths as |*σ_L_*| increases beyond a threshold greater than 1 as represented by the increase in (black plot is for *r_S_*/*r_L_* = 1/3; blue and green plots are for *r_S_*/*r_L_* = 1/2; in the green plot, *l, r_L_*, and *r_S_* have been rescaled by a factor of 1.5 compared with the blue plot), (b) Colour symmetry break. In all plots *σ_L_* = 2. As either *β* or *β_L_* increases, gyrating labyrinths bifurcate to uniform block colour attractors.

For colour symmetry (*β*,*β_L_* = 0), as depicted in Figures 3a and Movie S6, stationary labyrinths bifurcate to continually gyrating labyrinths as |*σ_L_*| increases beyond approximately 1.5 (*r_S_*/*r_L_* = 1/2) or 2.0 (*r_S_*/*r_L_* = 1/3). This is quantitatively represented in the plots of Figure 3a by the summary statistic *f_L_* which represents the time-averaged speed of movement of the interface, see Supplementary Material for details. *f_L_* appears to be unaltered when all length-scales are simultaneously rescaled while other parameters are held constant (compare the blue and green curves in the plot of Figure 3a), indicating that the structure of the bifurcation diagram is unaffected by discretisation effects of the lattice. For *σ_L_* = 2, the value of *r_S_*/*r_L_* must exceed a threshold of approximately 1/3 in order for this bifurcation to occur (not shown). An argument precisely analogous to that in the previous section explains why the bifurcation point for |*σ_L_*| must be greater than 1.

When colour symmetry is broken by perturbing *β* ≠ 0, gyrating labyrinths first freeze to be stationary labyrinths. They then bifurcate discontinuously to uniform block colour attractors once the magnitude of *β* exceeds a threshold as depicted in the upper plot of Figure 3b (*σ_L_* = 2 and *r_S_*/*r_L_* = 1/2). The bifurcation is similar when perturbing *β_L_* ≠ 0 but appears to be continuous.

### 3.3 Multi-colour hexagonal lattices and labyrinths from multi-colour lateral activation and inhibition

The 2 colour models in the previous sections are extended to an arbitrary number of colours denoted by *C_i,i_* = 1, 2, …, *n*. All symmetry breaking terms are omitted in this section. The reaction kinetics can then be described by a single expression:

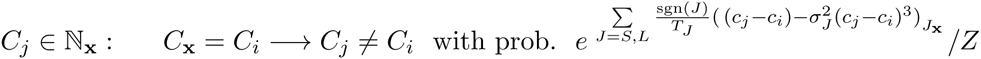

where

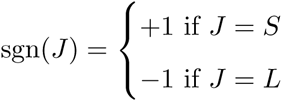

gives short-range activation and long-range inhibition, and 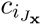 is the density of colour *C_i_* within range *r_J_*. We simulated 3 and 4 colours with activation at the interface and 1. long-range inhibition only; 2. short-range activation only; 3. both long-range inhibition and short-range activation; and 4. symmetry preserving nonlinear terms that compete with the short-range activation (|*σ_S_*| > 1) or the long-range inhibition (|*σ_L_*| > 1). The dynamics are portrayed in Figure 4.

**Figure 4:**
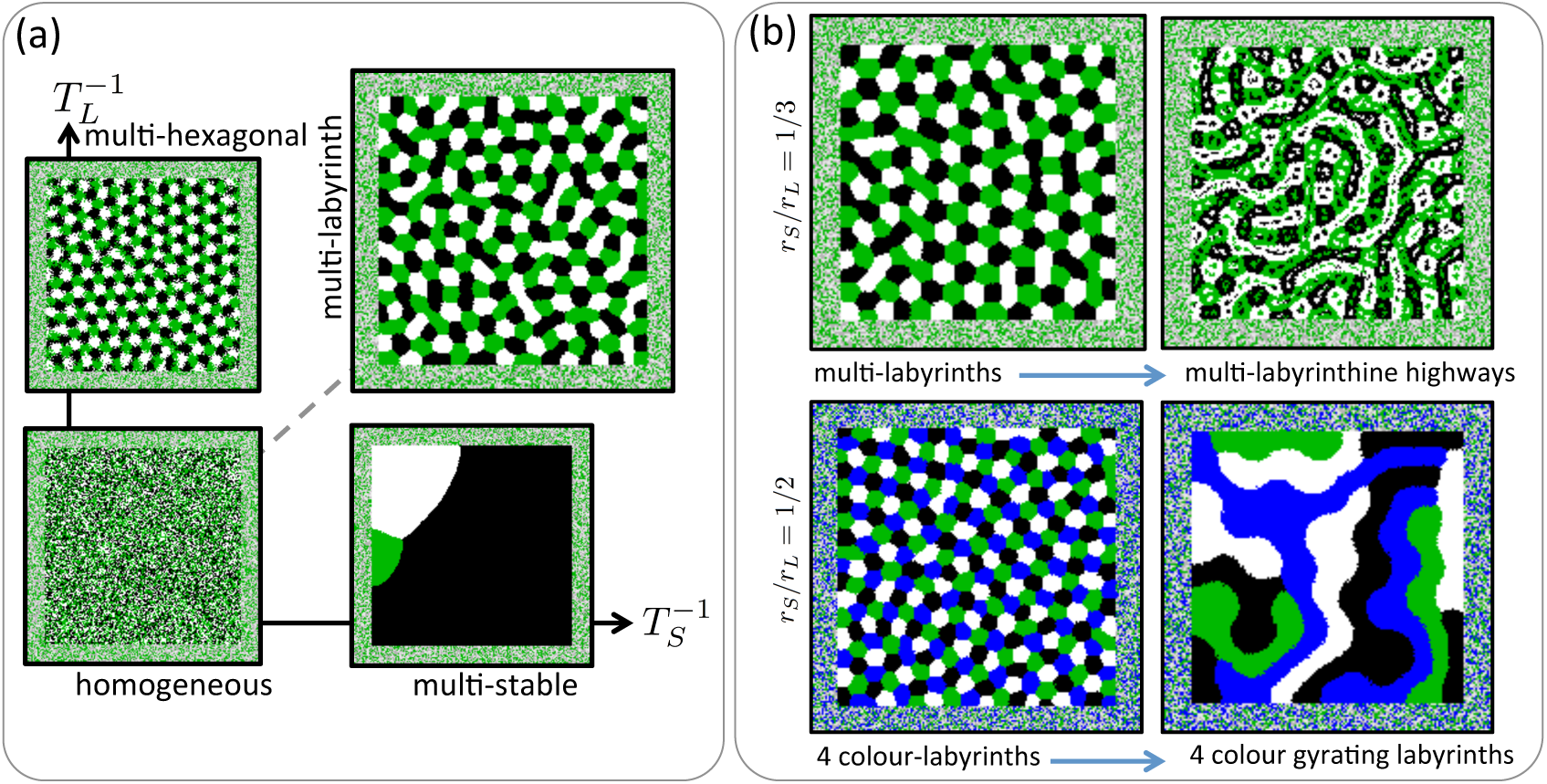
Patterning modes for multi-colour lateral activation and inhibition. (a) For no lateral feedbacks (*T_S_* = *T_L_* = ∞), attractors are homogeneous. For lateral inhibition only (*T_S_* = ∞, *T_L_* ≪ 1), attractors are a stationary multi-colour lattice, whereas for lateral activation only (*T_S_* ≪ 1, *T_L_* = ∞), attractors are multi-stable uniform block colour. Short-range activation and long-range inhibition together (*T_S_* ≈ *T_L_* ≪ 1) generate multi-colour labyrinths, (b) Increasing the nonlinear short-range inhibition |*σ_S_*| > 1 produces multi-colour labyrinthine high-ways (*r_S_*/*r_L_* = 1/3). Increasing the nonlinear long-range activation |*σ_L_*| > 1 produces gyrating labyrinths for 4 colours but for 3 colours attractors appear to remain stationary (*r_S_*/*r_L_* = 1/2).

As depicted in Figure 4a, lateral inhibition only (*T_S_* = ∞, *T_L_* ≪ 1) generates stationary hexagonal lattices. Lateral activation only (*T_S_* ≪ 1,*T_L_* = ∞) generates multi-stable block colour attractors where finally each site has the same colour and all of the permitted colours are equally likely. Short-range activation and long-range inhibition together (*T_S_* ≈ *T_L_* ≪1) generate multi-colour labyrinths. These attractors are analogous to 2 colour stripes, bistable uniform blocks, and labyrinths depicted in Figure 1.

When multi-colour labyrinths are perturbed by increasing the short-range symmetry preserving nonlinear competition |*σ_S_*| > 1, multi-colour labyrinths bifurcate to patterning modes that are analogous to labyrinthine highways for 3 and 4 colours, see Figure 4b. However, increasing the long-range symmetry preserving nonlinear competition |*σ_L_*| > 1 causes only 4-colour labyrinths to bifurcate to gyrating 4-colour labyrinths whereas 3-colour labyrinths bifurcate to attractors that appear not to gyrate.

### 3.4 Travelling stripes and spirals and reorganising labyrinths from cyclic symmetry breaking

When the model is extended to more than 2 colours, a new mode of symmetry breaking is possible which, unlike in Sections 3.1 and 3.2, does not enforce an accumulation of one particular colour. As in Section 3.3, because there are no symmetry breaking parameters that are associated with particular colours, the reaction kinetics can be described by a single expression:

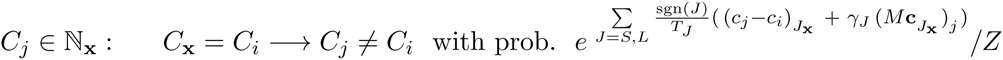

where again

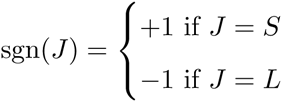

enforces short-range activation and long-range inhibition, and where 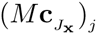 is the jth element of the vector

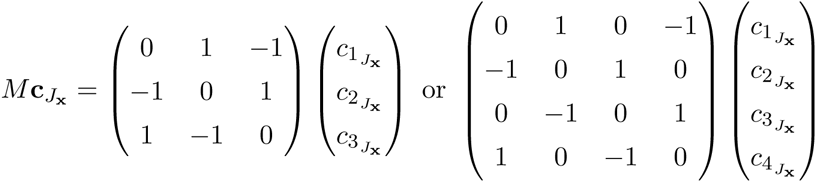

for 3 colours or 4 colours respectively, and *M* can be defined similarly for *n* colours. The circulancy of *M* drives the dynamics in a symmetry breaking cyclic colour ordering *C*_1_ → *C*_2_ → ⋯ → *C_n_* → *C*_1_ for *γ_S_*, *γ_L_* > 0. Colours can no longer be arbitrarily exchanged for one another without changing the model’s specification, yet there is no *a priori* propensity for one particular colour to accumulate. Reversing the sign of *γ_S_* (or *γ_L_*) drives the cycle in the opposite direction on the short (or long) range. A similar mode of symmetry breaking has been called cyclic dominance and studied in a spatial and probabilistic context within [16].

For lateral inhibition only (*T_S_* = ∞, *T_L_* ≪ 1), cyclic symmetry breaking with |*γ_L_*| ≈ 1 causes stationary multi-colour lattices to bifurcate to travelling stripes (3 colours) or travelling lattices (4 colours), see Figure 5 and Movie S7. For lateral activation only (*T_S_* ≪ 1, *T_L_* = ∞), cyclic symmetry breaking with |γ*_S_*| ≈ 1 causes multi-stable uniform block attractors to bifurcate to cyclic spiralling attractors, see Figure 5 and Movie S8. Strikingly, the focal points of the spirals appear to be stuck rigidly in one place and rarely wander within the domain. Simulations indicate that the number of spiral foci at the end of the simulation *N* varies linearly with 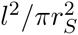, see Figure 5 inset panel, top plot; *N* appears to be unaffected by the simulation’s initial condition, see Figure 5 inset panel, middle and bottom plots.

**Figure 5:**
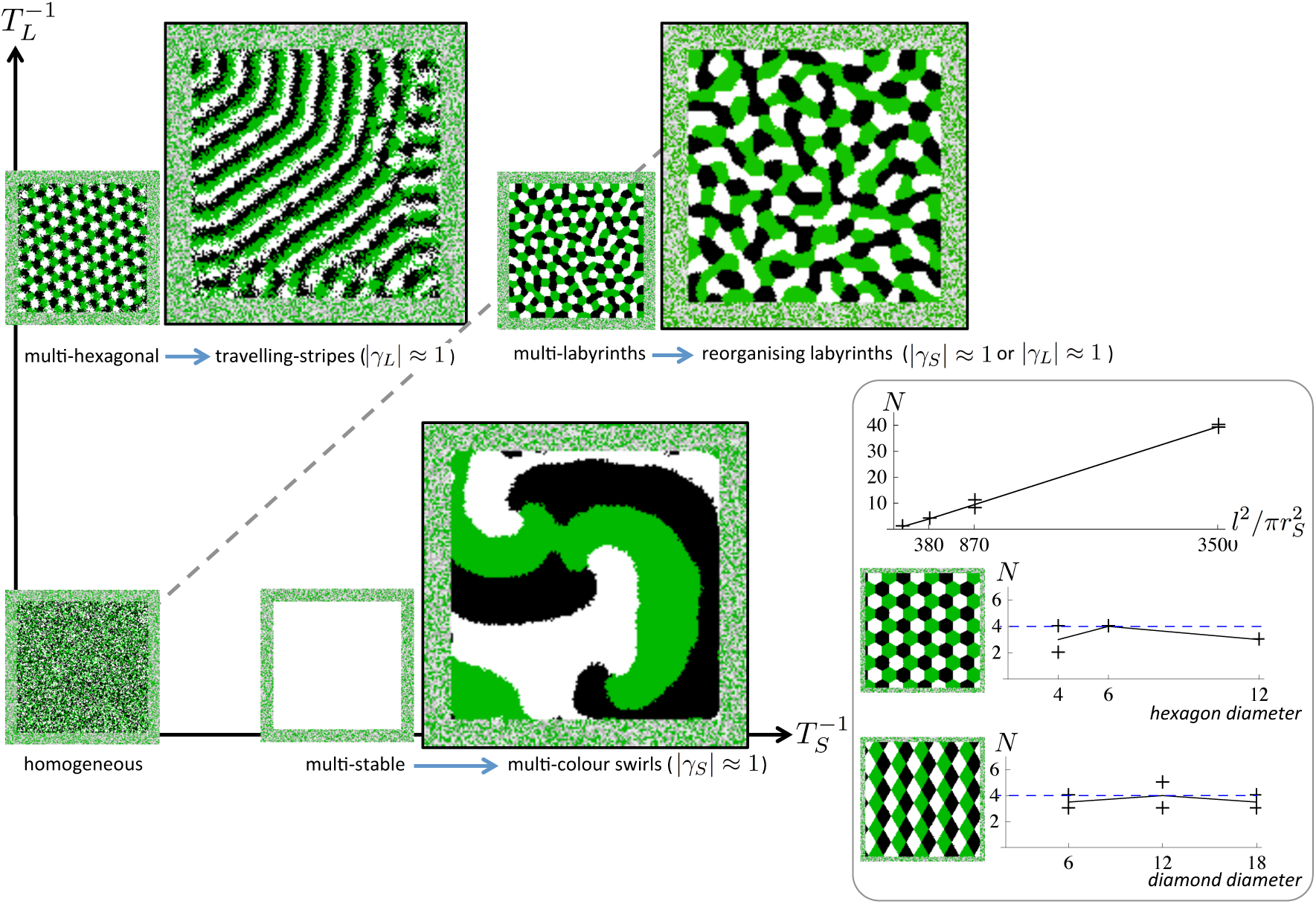
Travelling stripes, multi-colour swirls, and reorganising labyrinths from, cyclic symmetry breaking. For lateral inhibition only (*T_S_* = ∞, *T_L_* ≪ 1), increasing |*γ*_L_| ≈ 1 causes stationary multi-colour lattices to transit to travelling stripes/lattices (3 colours/4 colours); whereas for lateral activation only (*T_S_* ≪1, *T_L_* = ∞), increasing |*γ_S_*| ≈ 1 causes multi-stable blocks to transit to cyclic spirals. For both lateral activation and inhibition (*T_S_* = *T_L_* ≪ 1), increasing either |*γ_S_*| ≈ 1 or |*γ_L_*| ≈ 1 causes stationary multi-colour labyrinths to transit to dynamic, continually reorganising attractors. Panel: For cyclic spirals (*T_S_* ≪ 1, *T_L_* = ∞, |*γ_S_*| ≈ 1), the total number of spiral foci in the domain *N* varies linearly with 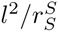 (top plot); changing the initial conditions to hexagonal/diamond lattices then varying the diameter of initial hexagons/diamonds (middle/bottom plot) appears to have little effect on *N* compared with the default initial condition (dashed line).

For short-range activation and long-range inhibition together (*T_S_* ≈ *T_L_* ≪ 1), cyclic symmetry breaking on the short or long range (|*γ_S_*| ≈ 1 or |*γ_L_*| ≈ 1) causes multi-colour labyrinths to continually reorganise, see Movies S9 and S10.

**Table 1:** Parameters for panels and plots in figures. The ‘end time’ is the number of transitions per simulation. *n* is the number of instances simulated for generating statistics. ‘t’, ‘m’, ‘b’, ‘p’ stand for ‘top row’, ‘middle row’, ‘bottom row’, ‘panel’ respectively in the corresponding figure or plot. ‘*V*’ stands for ‘varying’ in the corresponding figure or plot.

| Figure | $l$ | $\beta$ | $(r_S, T_S^{-1}, \beta_S, \sigma_S, \gamma_S)$ | $(r_L, T_L^{-1}, \beta_L, \sigma_L, \gamma_L)$ | end time | $n$ |
| --- | --- | --- | --- | --- | --- | --- |
| 1(c) t | 140 | $V$ | $(4, 0, -, -, -)$ | $(12, 16, -, -, -)$ | $3 \times 10^6$ | 5 |
| 1(c) b | 140 | $V$ | $(4, 16, -, -, -)$ | $(12, 16, -, -, -)$ | $3 \times 10^6$ | 5 |
| 2(b) t black/blue | 140/210 | 0 | $(4/6, 16, 0, V, -)$ | $(12/18, 16, -, -, -)$ | $2 \times 10^6$ | 5 |
| 2(b) b | 140 | 0 | $(V, 16, 0, 2, -)$ | $(12, 16, -, -, -)$ | $2 \times 10^6$ | 5 |
| 2(c) t black/blue | 140 | $V$ | $(4/6, 16, 0, 2, -)$ | $(12, 16, -, -, -)$ | $2 \times 10^6$ | 2 |
| 2(c) b black/blue | 140 | 0 | $(4/6, 16, V, 2, -)$ | $(12, 16, -, -, -)$ | $2 \times 10^6$ | 2 |
| 3(b) t black/blue | 140 | 0 | $(4/6, 16, 0, 0, -)$ | $(12, 16, 0, V, -)$ | $5 \times 10^6$ | 5 |
| 3(b) t green | 210 | 0 | $(9, 16, 0, 0, -)$ | $(18, 16, 0, V, -)$ | $5 \times 10^6$ | 5 |
| 3(b) b | 140 | 0 | $(V, 16, 0, 0, -)$ | $(12, 16, 0, 2, -)$ | $5 \times 10^6$ | 5 |
| 3(c) t | 140 | $V$ | $(6, 16, 0, 0, -)$ | $(12, 16, 0, 2, -)$ | $5 \times 10^6$ | 2 |
| 3(c) b | 140 | 0 | $(6, 16, 0, 0, -)$ | $(12, 16, V, 2, -)$ | $5 \times 10^6$ | 2 |
| 4(a) | 140 | – | $(4, 0/16/16, -, -, 0)$ | $(12, 16/0/16, -, -, -)$ | $3 \times 10^6$ | – |
| 4(b) | 140 | – | $(4/6, 16, -, 2/0, -)$ | $(12, 16, -, 0/3.4, -)$ | $3 \times 10^6$ | – |
| 5 | 140 | – | $(4, 0/16/16, -, -, 0/1/0)$ | $(12, 16/0/16, -, -, 1/0/2)$ | $3 \times 10^6$ | – |
| 5 p t/b | 210 | – | $(V/6, 16, -, -, 1)$ | $(-, -, -, -, -)$ | $3 \times 10^6$ | 2 |

## 4 Discussion

The models we have presented are simple generalisations of the well known pattern-producing dynamics of lateral activation and inhibition. Yet, despite their simplicity, to our surprise they have produced multiple examples of what are, to the best of our knowledge, qualitatively new dynamics and attractors including labyrinthine highways, Kagome lattices, gyrating labyrinths, and corresponding multi-species analogies. We anticipate that these attractors are robust to changes in details of the model such as the square geometry of the lattice or the precise formulation of the reaction kinetics and rather that they depend on the class of interactions implemented by the model. Whether the geometry of the square lattice impacts upon results could be better established by adapting our code, that is available on request and uses plugins from the software Processing [17], to run on hexagonal or Voronoi grids or by completing the derivation of the mean-field equations (see Supplementary Material) and simulating the resulting system of PDEs. We further anticipate that for each class of interactions all qualitatively new attractors were identified: a simple mean-field equation governing the model’s output allowed us to search for new patterns systematically.

**Table 2:**
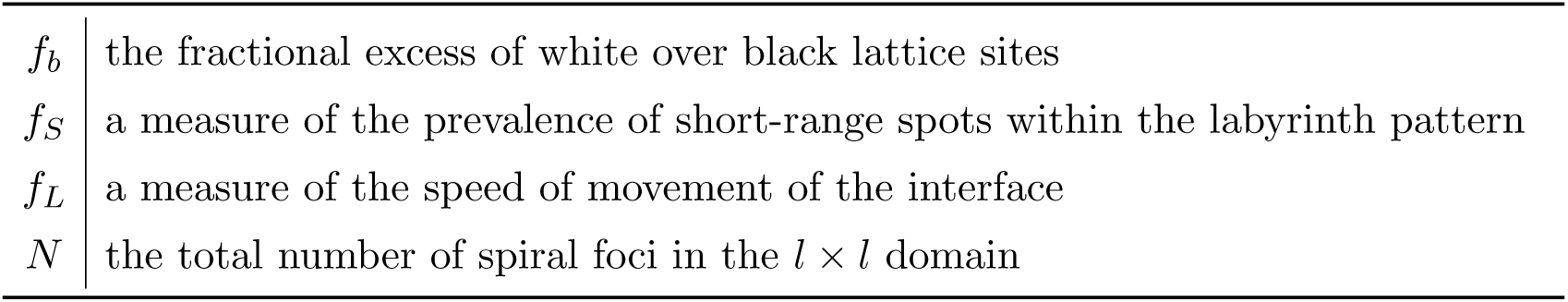
Definitions of summary statistics.

Since we believe that the model outputs do not depend on details of the implementation, we explore and speculate about the likely applicability of the models to biology. Firstly, our study indicates that new patterns—beyond stripes, labyrinths, and hexagonal nets—such as labyrinthine highways or Kagome lattices may be generated by a collection of cells with the following properties: i) a master regulator that once expressed initiates the expression of two diffusing ligands and a membrane-junction signal; ii) receiving the membrane-junction signal from a neighbouring cell is an absolute requirement for the expression of the master regulator; (iii) the first ligand diffuses in excess of a few cell lengths on average before being degraded; or it binds to cell-surface receptors to trigger a signal that decreases the likelihood of expression of the master regulator; (iv) the second diffusing ligand, which tends to be degraded within a range that is short compared with the range of the first diffusing ligand, upon binding to a receptor either increases or decreases the likelihood of expression of the master regulator depending on whether the number of bound receptors is below or above a threshold. However, it must be recognised that our model is only an abstract representation of i)–iv); in particular, it represents only cases where the time-scales of diffusion, degradation, and membrane junction signalling are much faster than the time-scale of the master regulator’s response to such signals. There are other possible long-range signalling mechanisms besides diffusion, operating over lengths which span multiple cells, that can account for lateral feedbacks: in animals, these include the active migration of cells, e.g. [18], and the dynamic protrusions of filopodia, e.g. [19, 20].

Secondly, our toy model motivates us to speculate about a possible dynamical feature of some developmental programs. In the model, when parameters enforce what we have called colour symmetry (*β* = 0)—a prerequisite for e.g. stripes and labyrinths—that region of parameter space is comparatively rich in the diversity of final patterns that are generated; consequently, the final pattern responds sensitively and flexibly to sustained changes in parameter values. The broad range of natural patterns suggests that this might be evolutionarily advantageous. While genetic drift would tend to take the system away from colour symmetry, we expect evolution to act as a tuning force which maintains the sensitivity and flexibility of developmental programs^1^.

## 5 Acknowledgements

This work began under the supervision of Tom Duke at the London Centre for Nanotechnology and CoMPLEX, University College London. Tom Duke is deeply missed. We thank David Wright for running a number of simulations and for stimulating discussions, Michael Cohen for introducing us to discrete models of lateral inhibition, and Pau Formosa-Jordan for a critique of the manuscript.

## Footnotes

1 We describe a dynamical system tuned to its critical point. A self-tuned critical system is known to operate in the inner ear where it affords tremendous sensitivity of response to vibrations of minute amplitude [21], but, as far as we are aware, no specific system is thought to operate in this manner owing to the action of evolution on the genome.

